# Fatty acid binding protein 5 forms higher order assemblies with FLAP or COX-2 in LPS-stimulated macrophages

**DOI:** 10.1101/2020.10.26.355594

**Authors:** Shelby E. Elder, Nicholas C. Bauer, Roy J. Soberman, Angela B. Schmider

## Abstract

Immune cells must integrate multiple extracellular signals to produce an appropriate inflammatory response, including production of a dynamic mix of eicosanoids and related bioactive lipids. Synthesis of these lipids is initiated on the membrane surface of the nuclear envelope and endoplasmic reticulum. One critical question is how the precursor arachidonic acid (AA) is distributed between the initial biosynthetic enzymes of the arachidonate 5-lipoxygenase (5-LO) and prostaglandin-endoperoxidase synthase-1/2 (COX-1/2) related pathways. To understand these balancing mechanisms, we hypothesized that fatty acid binding proteins mediate this process. We employed a multi-modal imaging approach by combining direct stochastic optical reconstruction microscopy (dSTORM) with computational analyses and fluorescence lifetime imaging microscopy (FLIM) to delineate the relationships of fatty acid binding proteins 3, 4, and 5 (FABP3–5) with 5-LO activating protein (FLAP), COX-1, and COX-2 in the presence of a stimulus (lipopolysaccharide, LPS) that triggers the synthesis of prostaglandin E_2_ (PGE_2_). LPS triggers a redistribution of FABP5 to higher order assemblies of COX-2 or FLAP. This was evidenced by a decrease in lifetime determined by FLIM. Colocalization between FABP3 and FLAP decreased, but no other changes in distribution were observed for FABP3 and FABP4. In contrast, assemblies of FABP5 with COX-1 were smaller and showed an increase in lifetime. The data indicate that FABP5 is a member of higher order assemblies of eicosanoid biosynthetic enzymes and that FABP5 may play a key role in regulating the organization of these structures. FABP5 is positioned to distribute AA to both the 5-LO and COX-2 pathways.

## Introduction

How cells integrate extracellular signals to give an appropriately balanced response is a problem shared by all signaling pathways. In the case of eicosanoids and related bioactive lipids, cells must generate a balanced and shifting mix of products depending on the extracellular environment and strength and variety of signaling inputs. The membrane surfaces of the nuclear envelope and the ER are where synthesis is organized and initiated. One critical issue is how the distribution of arachidonic acid (AA) is dynamically parsed between the initial biosynthetic proteins of the 5-Lipoxygenase (5-LO) pathway, or to the Prostaglandin-endoperoxide synthase-1 (PTGS1, COX-1) or PTGS2 (COX-2) related pathways, which are key potential regulatory junctions.

Fatty acid-binding proteins (FABPs) are a family of nine small (13-14 kDa) proteins which are highly expressed in the cytosol. They function by binding a variety of hydrophobic ligands (1–4) and transporting them to integral membrane proteins on the interfacial surface of membranes (4). Due to their role as chaperones that distribute intracellular lipids, FABPs can function as regulators of inflammatory pathways and eicosanoid production (5, 6). There are ten different FABP isoforms (FABPs 1-7, 8 (Myelin P protein), 9 and 12) that are expressed in diverse cells and tissues including macrophages, dendritic cells, liver, heart, brain and kidney (7). Most FABPs can bind to AA with moderate or strong affinity (FABPs 1, −2, −3, −5, −7, −8) (8). The binding of AA to FABP4 and FABP5 alters their conformation and exposes a nuclear localization signal (9, 10). Both FABP4 and FABP5 translocate to the nucleus and activate members of the peroxisome proliferator-activated receptor family by delivering lipid agonists (2, 9, 11–14). FABPs increase the half-life of LTA_4_, the precursor of other leukotrienes (LTs) (5). While interaction of FABPs with peroxisome proliferator-activated receptors (PPARs) and stabilization of LTA_4_ are established, FABP3, FABP4, and FABP5 are all capable of binding AA, and FABP5 has been shown to preferentially bind AA over LTA_4_ (15, 16). Bogdan et al. demonstrated that FABP5 impacts the production of prostaglandin E_2_ (PGE_2_) by induction of mPGES-1 (which converts the COX-1/2 product PGH_2_) through NF-κB, and also functioning as a shuttle of AA (17). Thus, each of these FABPs have the potential to modulate the earliest steps in the synthesis of PGs and LTs.

To understand how cells use intracellular membranes as platforms to organize cell signaling we tested the hypothesis that FABPs could be important structural and functional members of higher order assemblies of eicosanoid biosynthetic enzymes. We focused our experiments on FABP3, which is ubiquitously expressed in mammalian cells, and FABP4 and −5, which are highly expressed in macrophages (2). We employed a multi-modal imaging approach by combining direct stochastic optical reconstruction microscopy (dSTORM) with computational analyses and Fluorescence Lifetime Imaging Microscopy (FLIM) to delineate the relationships of FABPs 5, 4, and 3 with FLAP, COX-1, and COX-2 in response to a stimulus that triggers the synthesis of PGE_2_. LPS triggers a redistribution of FABP5, but not FABP4 or FABP3 to higher order assemblies of COX-2 or FLAP on the nuclear envelope and ER. Furthermore, a decrease in τ_1_ between FABP5 and either FLAP or COX-2 was found. In contrast, the assemblies of FABP5 with COX-1 are decreased in size and show an increase in τ_1_. The data support the concept that FABPs are previously unrecognized constituents of the higher order membrane assemblies of enzymes and proteins in eicosanoid synthesis. The incorporation of FABPs is likely to be a key regulatory step in organizing the initiation of inflammatory signaling.

## Results

### FABP3, −4 and −5 are expressed in RAW264.7 cells

After 18 h stimulation with lipopolysaccharide (LPS), the supernatant was removed from RAW264.7 cell cultures and analyzed for PGE_2_. Stimulation with LPS resulted in 30-fold increase over controls (**Figure 1A**). RNA was then extracted from the cells and screened by RT-PCR using PCR primers for FABPs 1-7, −9 and −12 (**Table S1**). We only detected expression of FABP3, −4 and −5 (**Figure 1B**). Western blots with antibodies for FABP3/4/5 were used to confirm the expression of the proteins in activated cells. We observed strong bands for each protein, running slightly larger than the expected size (~15 kDa) (**Figure 1C**). FABP4 and FABP5 blots also show a second band at approximately twice the size. Because FABPs form homodimers, the most likely explanation is that a small fraction can remain associated.

**Figure 1.**
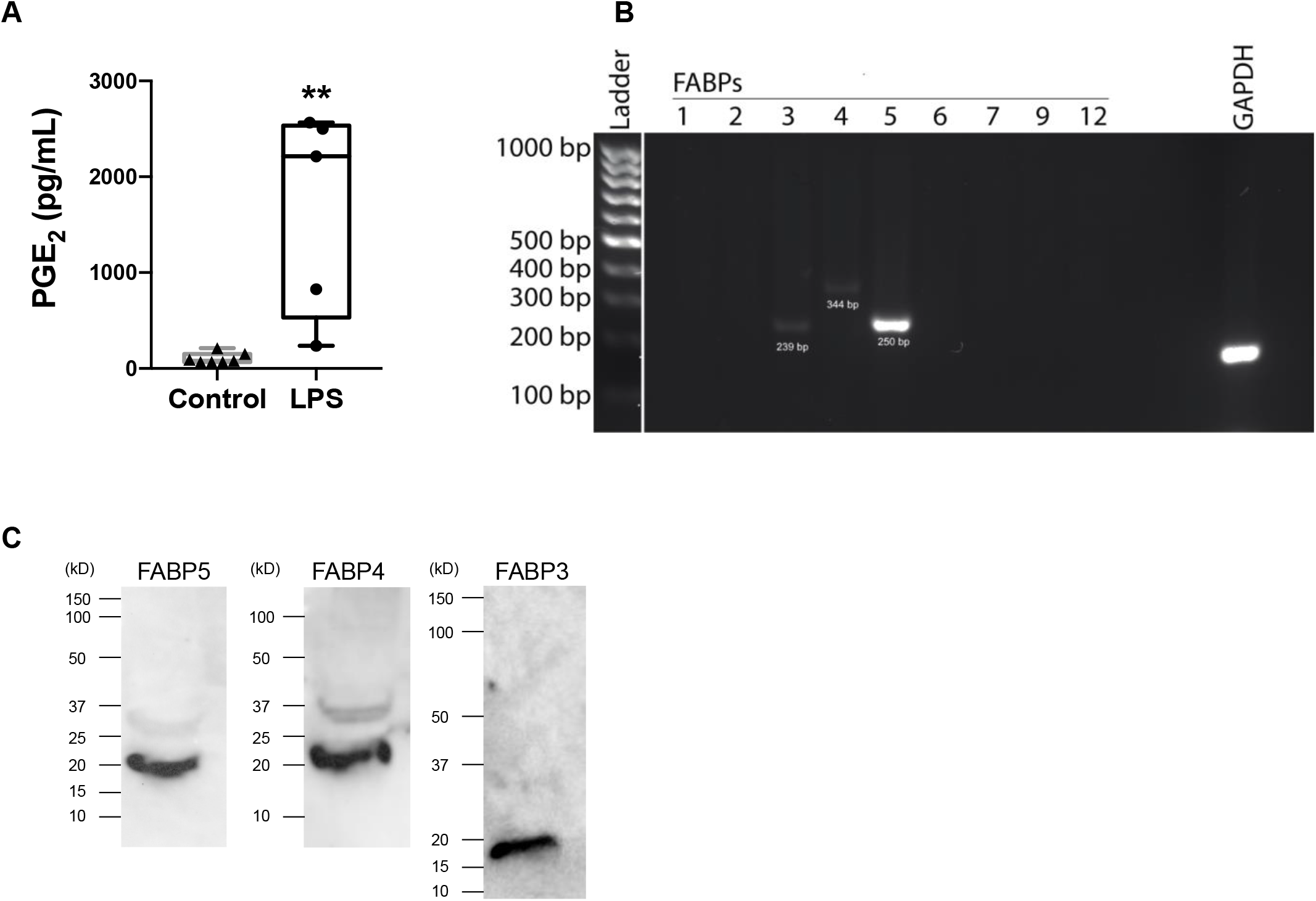
FABP3, −4, and −5 are expressed in RAW cells. (A) PGE_2_ (pg/mL) in cell supernatants generated in response to LPS (100 ng/mL) for 18 hours in RAW cells (300,000 cells/well). (B) FABP isoforms 1-7, 9, and 12 were analyzed. RT-PCR identified FABP3, −4, and −5 to be the isoforms expressed in RAW cells. Equal amounts of cDNA were loaded to each well. cDNA bands at 239 bp, 344 bp, and 250 bp, identify FABP3, −4, and −5 respectively. GAPDH was used as a loading control. (C) Western blot for protein expression of FABP3, −4, and −5 in RAW cell protein extract. 30 μg of protein was loaded in each lane. Student’s *t*-test was performed where ** = p < 0.005. The boxes show the median and interquartile range. Three separate experiments were used.

### LPS selectively re-organizes the associations of FABP5 in cells

**Figure 2** shows analysis of the relationships of FLAP with FABPs 5, 4, and 3 determined by the Clus-DoC algorithm. RAW cells were stimulated with LPS for 18 hours and the media removed and assayed for PGE_2_ to confirm activation (**Figure 1A**) and analyzed by Clus-DoC. Stimulated and control cells were washed, fixed, and analyzed by two-color dSTORM imaging (**Figure S1**). Two-color dSTORM analysis was performed for FABPs 3, 4, and 5, and FLAP (**Figure 2**). PCR screens for all other FABPs showed no bands. The relationships of the two proteins were analyzed using Clus-DoC; DoC scores ranging from −1 (anti-correlated) to 1 (correlated) were determined. Localizations with a DoC score ≥ 0.4 were considered colocalized. **Figure 2A** shows that in unstimulated cells, the integral membrane protein FLAP (green) is localized to the nuclear envelope and also distributed through the endoplasmic reticulum (ER). After stimulation with LPS, there is a clear shift to localizations with a DoC score greater than 0.4 as shown on both the histogram and the co-localization map, indicating a clear increase in co-localized molecules. The localization and co-localization graphs support the observation that there is greater than a three-fold increase in the co-localization of FABP5 and FLAP in the nuclear envelope (0.17 [0.10, 0.23] to 0.57 [0.43, 0.72]) (**Figure 2A**). In contrast, there was no effective change in the relationship of FABP4 and FLAP in the response to LPS (**Figure 2B**). FABP3 showed a slightly decreased co-localization with FLAP (**Figure 2C**).

**Figure 2.**
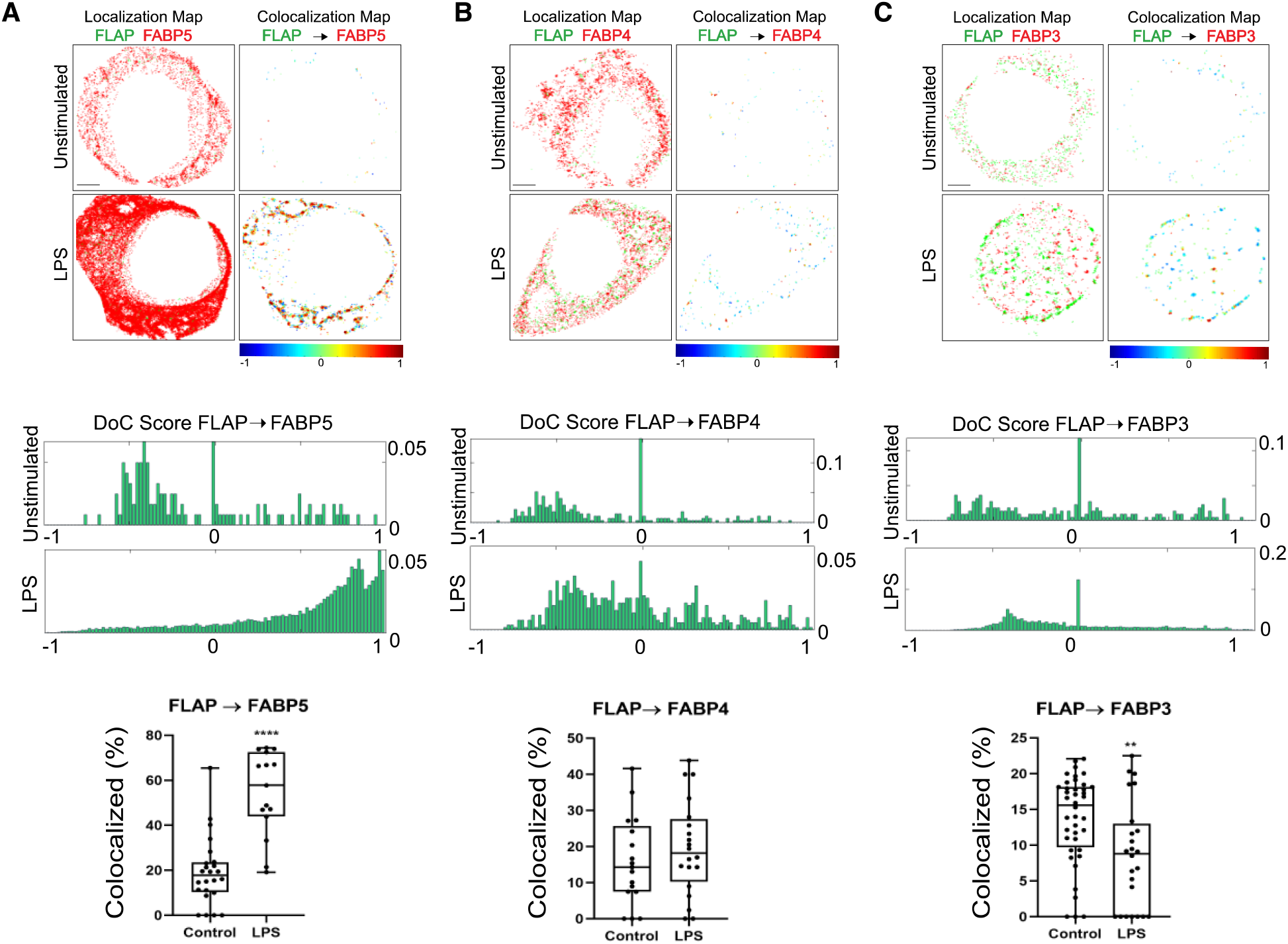
The relationships of FLAP with FABPs. (A) FABP5, (B) FABP4, and (C) FABP3. 2-color d-STORM was used to localize FLAP and FABPs in the absence and presence of 100 ng/mL LPS. To determine the relationships between FABPs and FLAP, Clus-DoC was utilized to analyze localizations and generate both histograms and colocalization maps. Histograms of DoC scores for all molecules FLAP (green) are shown from representative cells, and the co-localization scores were used to generate the scale shown below the co-localization map. The change in percent of FLAP co-localization was determined. Statistical significance was assessed by Student’s *t-test* with significance indicated by ** p < 0.005, **** p < 0.0001. The boxes show the median and interquartile range. From 4 to 21 cells over 3 separate experiments. Scale bar 2.0 μm.

When the relationships of COX-2 to FABPs 3, 4, and 5 were analyzed, the results were similar to FLAP (**Figure 3**). After LPS stimulation, the percentage of COX-2 associated with FABP5 increased in percent from 19 [7, 33] to 31 [25, 47] whereas there was no percent change following LPS in the relationship of COX-2 with either FABP4 (21 [11, 48] to 18 [14, 33]) or FABP3 (17 [8, 26] to 31[16, 37]) (**Figures 3B and C**). We next examined the relationship between FABP3 and FABP4 with COX-1 (**Figure 4**). Prior to stimulation with LPS, 17% of COX-1 (17 [9, 40]) interacted with FABP5, but then decreased after LPS treatment (11 [6, 14]) (**Figure 4A**). There was no change in the relationship between COX-1 and FABP4 (19 [12, 39] to 21 [16, 33]) or FABP3 (17 [6, 24] to 21 [8, 24]) (**Figures 4B and C**).

**Figure 3.**
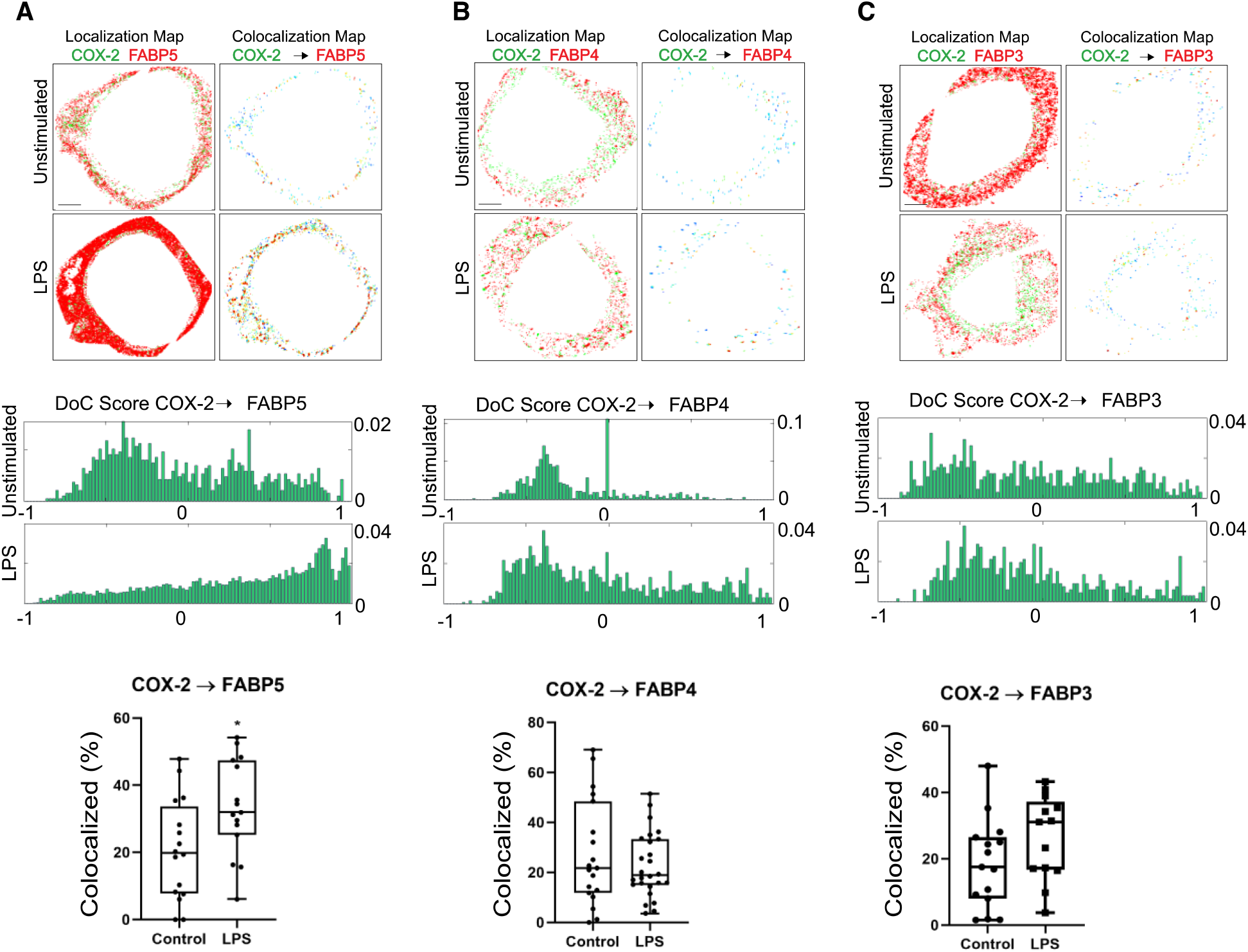
The relationships of COX-2 with FABPs. A) FABP5, (B) FABP4, and (C) FABP3. 2-color dSTORM was used to localize COX-2 and FABPs in the absence and presence of 100 ng/mL LPS. To determine the relationships between FABPs and COX-2 Clus-DoC was utilized to analyze localizations and generate both histograms and colocalization maps. Histograms of DoC scores for all molecules FLAP (green) are shown from representative cells, and the co-localization scores were used to generate the scale shown below the co-localization map. The change in percent of COX-2 co-localization was determined. Statistical significance was assessed by Student’s *t-test*, with significance indicated by * p < 0.05. The boxes show the median and interquartile range. From 4 to 21 cells over 3 separate experiments.

**Figure 4.**
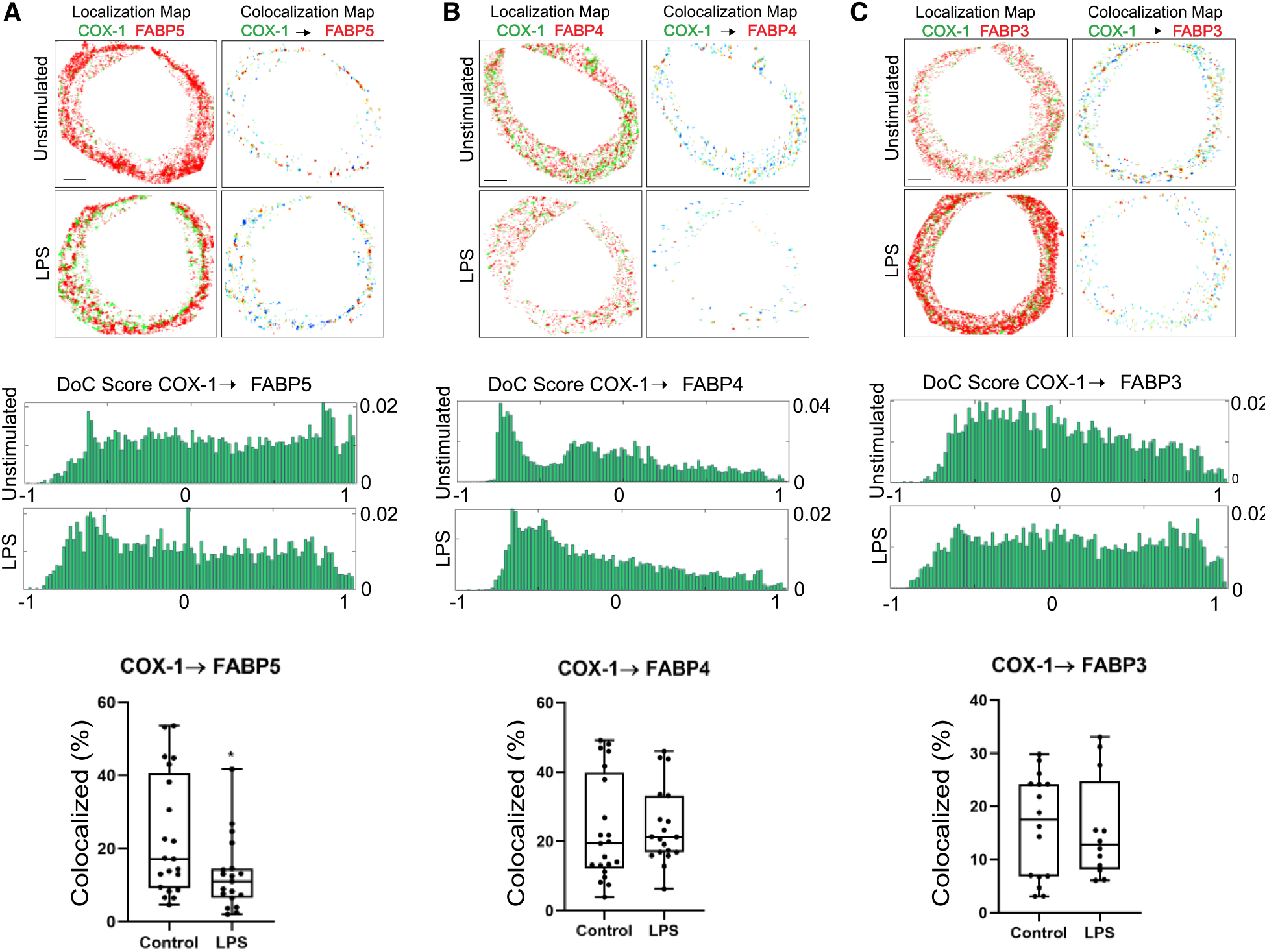
The relationships of COX-1 with FABPs. A) FABP5, (B) FABP4, and (C) FABP3. 2-color dSTORM was used to localize COX-1and FABPs in the absence and presence of 100 ng/mL LPS. To determine the relationships between FABPs and COX-1 Clus-DoC was utilized to analyze localizations and generate both histograms and colocalization maps. Histograms of DoC scores for all molecules COX-1 (green) are shown from representative cells, and the co-localization scores were used to generate the scale shown below the co-localization map. The change in percent of COX-1 co-localization was determined Statistical significance was assessed by Student’s *t-test*, with significance indicated by * p < 0.05. The boxes show the median and interquartile range. From 4 to 21 cells over 3 separate experiments. Scale bar 2.0 μm.

To characterize potential higher order assemblies, we employed additional features of Clus-DoC to analyze relationships FABP5:FLAP (Figure 5), FABP5:COX-2 or FABP5:COX-1 (Figure 6). Clusters were defined as having ≥ 5 localizations of FABPs, COX-1, or COX-2, or ≥10 FLAP localizations. We classified these clusters into Low Interacting Clusters (LIC, orange line) and High Interacting Clusters (HIC, gray bar with black line) which have at least 5 and 10 localizations, respectively, with a DoC score ≥ 0.4. Clusters with no interaction (NIC, white bar with black line) were also identified. Multiple changes in the relationship of FABP5:FLAP were identified (Figure 5) after stimulation with LPS. Because we previously observed strong clustered interactions of FLAP, 5-LO, and cPLA_2_ associated with the synthesis of LTs (18, 19) we focused on the changes we observed in HICs with FABP5. The largest increases between FABP5:FLAP following LPS exposure were in the cluster area of HIC, and the relative density of FLAP clusters (**Figure 5C and E**). The number of FLAP localizations in HIC clusters increased from 7 [5, 10]) to 13 [5, 44) and decreased in LIC clusters from 7 [4,8] to 2 [10.6, 4] following LPS (**Figure 5D**). The other HIC clustering parameters did not change following LPS stimulation (**Figures 5A, B and E**). Changes in NIC and LIC were observed following LPS treatment (**Figures 5B, C and E**). LPS caused no changes in FABP5 cluster relative density (**Figure 5F**). Relative density is calculated as the average of local relative density within 20 nm of each localization in the cluster, divided by the average cluster relative density, providing a measure of the distribution of localizations within a cluster. There were no major changes in the percent of COX-1 interacting in clusters, COX-1 relative density in clusters or FABP5 relative density in clusters with COX-1 in response to LPS (**Figure S2**). Similarly, in response to LPS, there were no observed COX-2 molecules interacting in clusters, average COX-2 cluster area, average number of COX-2 per cluster, COX-2 relative density in clusters or FABP5 in clusters with COX-2 (**Figure S3**). Overall, the average number of FLAP with FABP5 and FABP4 localizations per ROI increased following LPS exposure from 250 [119, 1142] to 4006 [2764, 6985] and 2979 [1366, 5388] to 5323 [2824, 10797] and 5719 [3404, 6841] to 9271 [6841, 14810], respectively (**Table S2**).

**Figure 5.**
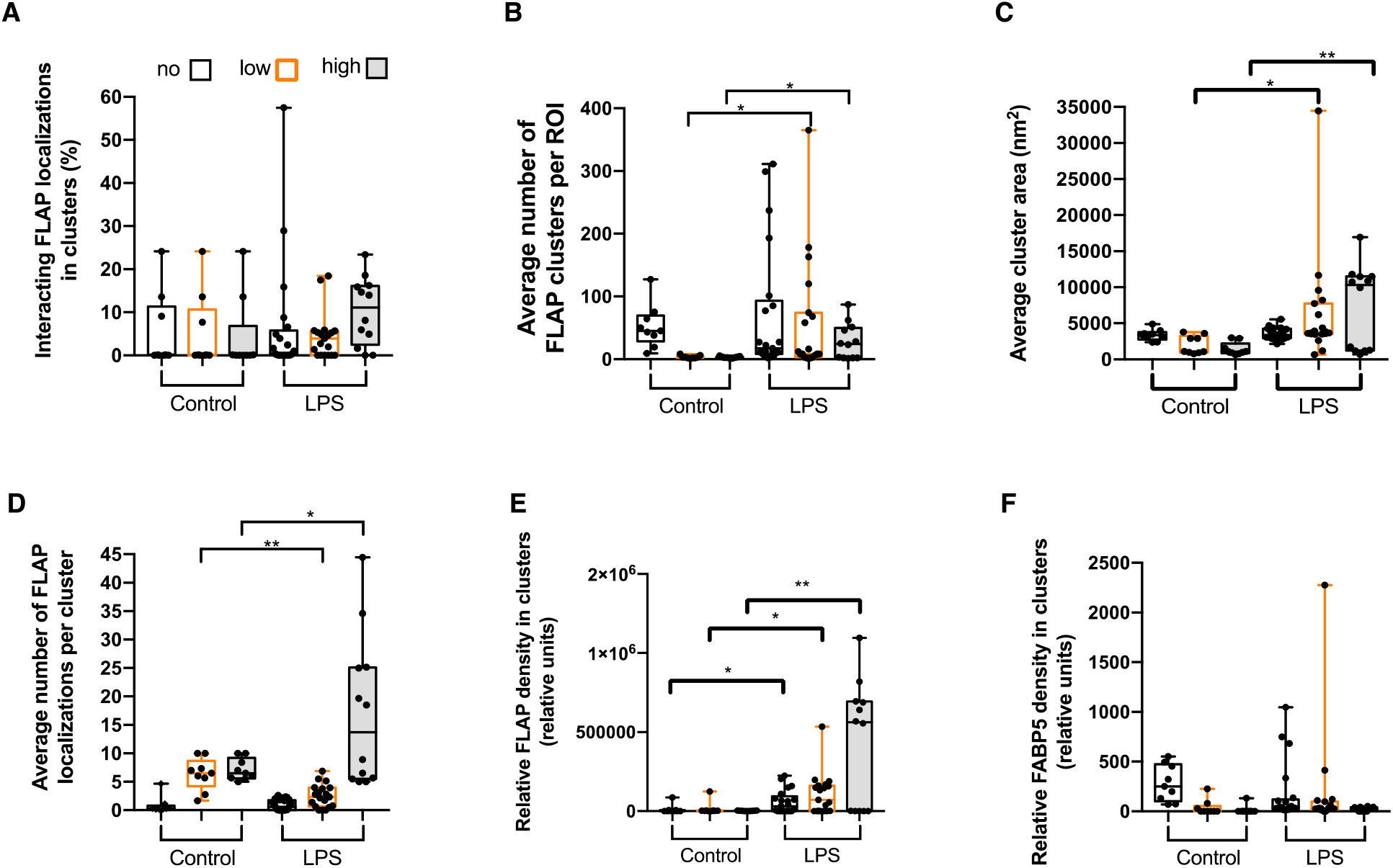
Activation with LPS reshapes FABP5 and FLAP co-cluster properties. Cells were treated with 100 ng/mL LPS for 18 hours or left untreated (control). Localization data was collected by two-color dSTORM and analyzed by Clus-DoC. Clusters were defined as having ≥ 5 FABP localizations and ≥10 FLAP localizations. These included Low Interacting Clusters (LIC, orange line); and High Interacting Clusters (HIC, black line with gray fill), which have a DoC score ≥ 0.4. Clusters with no interaction (NIC, black line) were also identified. (A) Percent of interacting FLAP localizations in clusters. (B) Average number of FLAP clusters per ROI. (C) Average clusters area. (D) Average number of FLAP localizations per cluster. (E) Relative FLAP relative density in clusters. (F) Relative FABP5 relative density in clusters. One-way ANOVA with Tukey’s post-hoc multiple comparisons test was performed to determine significance among interaction group where * p < 0.05 and ** p > 0.005. The boxes show the median and interquartile range. From 4 to 21 cells over 3 separate experiments.

**Figure 6.**
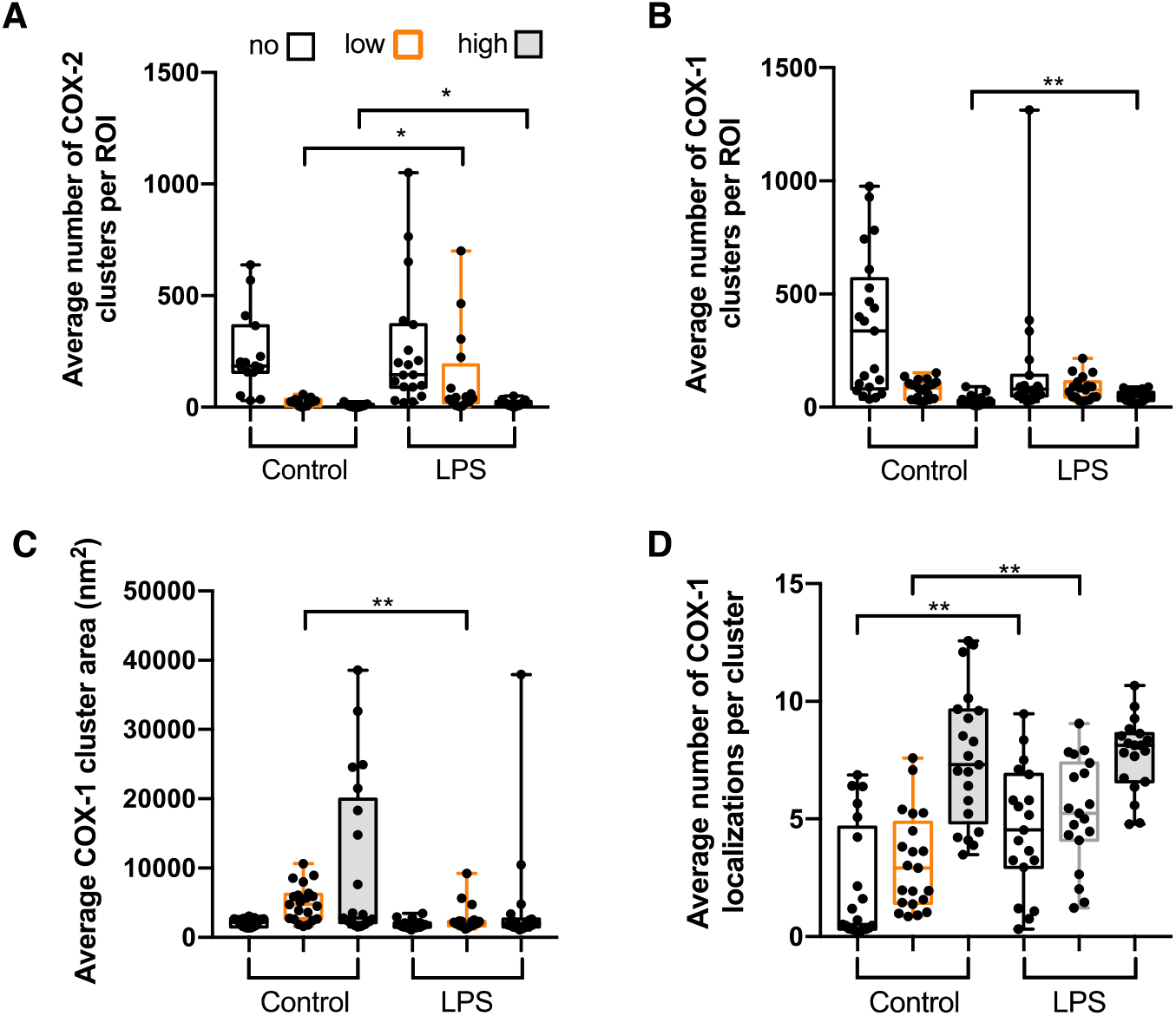
Changes in the cluster properties of FABP5 with COX-1 and COX-2 in response to LPS. Cells were treated with 100 ng/mL LPS for 18 hours or left untreated (control). Localization data was collected by two-color dSTORM and analyzed by Clus-DoC. Clusters were defined as having ≥ 5 FABP or COX-1/2 localizations and were split into three categories based on the number of localizations with a DoC score ≥ 0.4 indicating interaction where no interaction clusters (NIC, black line), low interaction clusters (LIC, orange line) having between 1 and 4 interacting localizations and high interaction clusters (HIC, black line with gray fill) having ≥ 5 interacting localizations. (A) Average number of COX-2 clusters per ROI. (B) Relationships of COX-1 with FABP5: Average cluster area; average number of localizations per cluster; Average number of clusters per ROI. One-way ANOVA with Tukey’s post-hoc multiple comparisons test was performed to determine significance among interaction groups where *p < 0.05; **p < 0.005. The boxes show the median and interquartile range. From 4 to 21 cells over 3 separate experiments.

We observed several differences between the FABP5:COX-2 and FABP5:COX-1 relationships. In the case of COX-2, the average number of HIC between FABP5:COX-2 increased from 5 [1, 13] to 13 [5, 26] after LPS stimulation and the number of LIC increased from 25 [6, 35] to 42 [20, 189] (**Figure 6A**). In the case of COX-1 (**Figure 6B**), only the average number of COX-1 HIC increased after LPS stimulation from 21 [11, 31] to 45 [29, 65] (**Figure 6B**). There were changes in the average COX-1 cluster area and average number of COX-1 localizations per cluster of NIC and LIC following LPS stimulation (**Figure 6C and D**). There was a significant overall increase in the average number of localizations per ROI following LPS stimulation for COX-2, 1281 [753, 2703] to 2995 [1874, 13884], but not for COX-1, 2912 [2189, 8006] to 5829 [2692, 7150] (Table S2).

The colocalization patterns observed using dSTORM were supported by FLIM analyses (**Table 1**). Unstimulated (control) and LPS-stimulated RAW cells were prepared and imaged using FLIM. FABP5 was designated as the donor fluorophore (Alexa Fluor 488-Affinipure donkey anti-rabbit IgG (H+L)) and FLAP, COX-1, or COX-2 was designated as the acceptor fluorophore (Alexa Fluor 594-Affinipure donkey anti-goat or anti-mouse IgG (H+L)). Significant decreases in donor lifetime (**τ**_1_) were observed for the FABP5:FLAP (2196 ± 50 ps to 1706 ± 48 ps). Interestingly, in response to LPS, the percent of interacting molecules (a_1_%) between FABP5:FLAP decreased from 28.51 to 2.5. Similarly, LPS caused a decrease in **τ**_**1**_ (2058 ± 46 ps to 1193 ± 77 ps) between FABP5:COX-2 and a decrease in a_1_% from 54 ± 94 to 27 ± 03. In the case of both protein pairs, these data imply that the quality of interaction is more critical than the quantity of interacting molecules and that higher order assembly of proteins is based on the quality of protein interactions. An additional factor is the large LPS-induced increase in abundance of FABP5 and COX-2 molecules, which could reduce the ratio of interacting molecules to total molecules without indicating a reduction in quantity of interactions. In contrast, **τ**_**1**_ increased in response to LPS for the FABP5:COX-1 pair (1700 ± 89 to 2245 ± 35 ps) with no change in a_1_% values. FABP5’s colocalization with COX-2, which is expressed in inflammatory response, rather than the constitutively expressed COX-1, suggests that this is an interaction unique to the LPS-induced inflammatory response. Moreover, we detected a decrease in **τ**_**1**_ between COX-2:FLAP in response to LPS (1315 ± 92 to 939 ± 62), suggesting that FABP5, COX-2 and FLAP co-associate to produce prostaglandins.

**Table 1.** τ_1_(ps) and a_1_(%) values measured by FLIM for FABP5:FLAP, FABP5:COX-2, and FABP5:COX-1 in unstimulated and LPS conditions. Students *t-test* was performed where *** = p < 0.001 and **** = p < 0.0001.

| Condition | Protein Pair<br>(Donor: Acceptor) | $\tau_1$(ps) | $a_1$(%) |
| --- | --- | --- | --- |
| Unstimulated | FAPB5: FLAP | 2196±50 | 48.15±2.5 |
| LPS | FAPB5: FLAP | 1706±48**** | 28.51±2.5**** |
| Unstimulated | FAPB5: COX-2 | 2058±46 | 54.94±3.3 |
| LPS | FAPB5: COX-2 | 1193±77**** | 27.03±3.0**** |
| Unstimulated | FAPB5: COX-1 | 1700±89 | 33.1±3.4 |
| LPS | FAPB5: COX-1 | 2245±35**** | 33.4±3.0 |
| Unstimulated | COX-2:FLAP | 1315±92 | 28±1.0 |
| LPS | COX-2:FLAP | 939±62*** | 27±1.0 |

## Discussion

The LPS-induced clustering of FABP5 with the integral membrane proteins initiating bioactive lipid synthesis are supported by two complementary imaging modalities. To detect reorganization of synthetic enzymes under conditions that only stimulate PGE_2_ generation but not LT formation, we utilized RAW cells stimulated by LPS. By focusing on the interactions of FLAP, COX-2, and COX-1 with FABP5 we were also able to probe the earliest steps segregating protein assemblies between these pathways. The regulation of the FABP5:FLAP and FABP5:COX-2 associations share common themes. These themes include not only increased co-localization, but a clear increase in the relative density of HIC. In both cases the FLIM data supported tighter associations in a subset of co-clustered molecules. In the case of COX-1, while the number of high interaction clusters (HIC) per ROI increased slightly, the **τ**_**1**_ also increased in the presence of an unchanged a1(%), showing no change in interaction. Thus, these results indicated a tight association between FABP5 with COX-2 and also FLAP, but not COX-1.

Prior studies have indicated that LPS primes neutrophils, macrophages, and eosinophils to generate large amounts of LTB_4_ or LTC_4_ after a second stimulus (20, 21). The explanation proposed was that LPS increased levels of cPLA_2_ which, in turn, increased the release of AA with the subsequent generation of more product (5, 22). Secretory PLA_2_ (sPLA_2,_ type V) also can supply AA to COX-2 and have crosstalk with cPLA_2_ in the initiation of eicosanoid production (23, 24). FABP5 facilitates transfer and modulates available concentrations of AA, thus the partitioning of FABP5, an interfacial lipid transport protein, could provide an additional regulatory mechanism that would collect and distribute AA substrate to COX-2 or FLAP. Since we show a close association of FAPB5 with FLAP and COX-2, direct protein-protein interactions could also regulate FLAP or COX-2 function independent of a substrate.

It is increasingly recognized that FABPs, and in particular FABP5, play roles in both assembling signaling complexes and simultaneously transferring substrates that play an important role in signaling. FABP5 links the function of fatty acid synthase (FASN) and monoacylglycerolipase (MAGL) to signaling by delivering lipid activators to nuclear hormone receptors including PPARδ and estrogen-related receptor α (ERRα) (25, 26). FABP5 is also responsible for the integrity of the mitochondria in lipid metabolism in regulatory T cells (27). These observations are suggestive that FABP5 has the potential to play both a shuttle and organizing role in the synthesis of bioactive lipids. Additionally, cPLA_2_ is localized with eicosanoid biosynthetic complexes in neutrophils and mast cells, so one possibility is that FABP5 functions to capture AA released by cPLA_2_ and link it to FLAP and COX-2. The results also suggest an inflammatory superassembly composed of FLAP, COX-2 and potentially LTC_4_ synthase, all integral membrane proteins. This would be brought together by AA released by cPLA2 and bound to FABP5 which would also serve in a structural role to assemble the complex. The next tier of enzymes, 5-LO and prostaglandin synthases, could then be recruited to their appropriate partners. However, we cannot rule out the possibility of other PLA_2_ enzymes, particularly sPLA_2_-V, and AA bound to FABP5, regulate AA as a substrate for eicosanoid production. This concept is depicted in **Figure 7**. Our work suggests that FABP5 could be a therapeutic target for reducing inflammation mediated by bioactive lipids. Small molecule inhibitors of FABP5 are being developed for prostate cancer and leukemia (28–30). These molecules and related FABP5 inhibitors should be tested for efficacy against inflammatory diseases.

**Figure 7.**
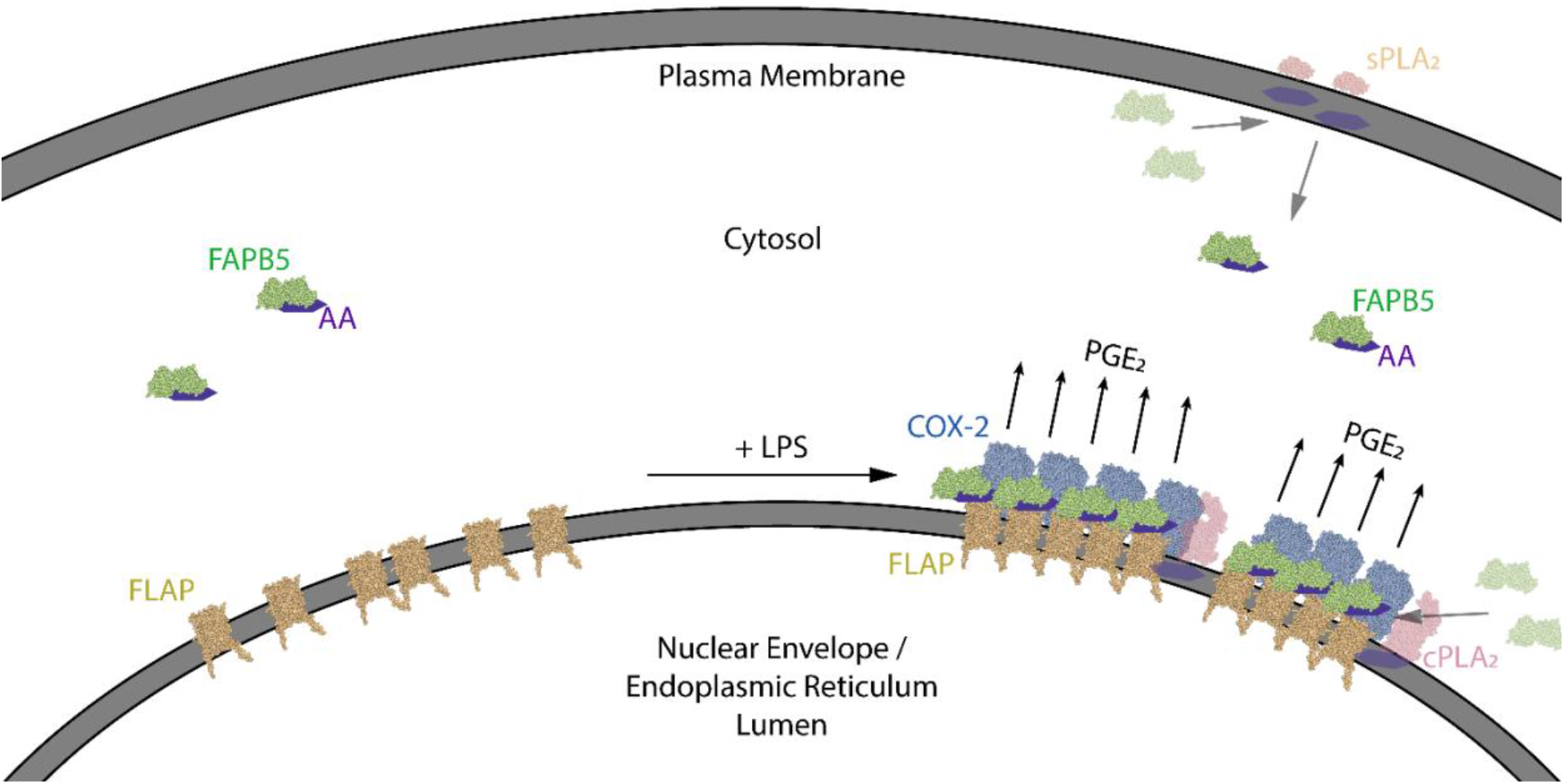
Proposed membrane organization of prostaglandin biosynthesis on membranes. The associations of the FABP5 (green), FLAP (gold), COX-2 (blue), and sPLA_2_/cPLA_2_ (faded orange/red, respectively) on the outer nuclear envelope and ER in unstimulated and LPS-activated macrophages. LPS increases levels of AA (purple hexgon), potentially released by cPLA_2_ or sPLA_2_, which are bound by FABP5, and induces clustering of FABP5 with FLAP and COX-2. COX-2 synthesizes prostaglandin H_2_, and downstream enzymes convert it to products like PGE_2_. Graphic was created using NGL Viewer (33) from RCSB PDB: COX-2 (6OFY), cPLA_2_ (1CJY), sPLA_2_ (1KQU), FABP5 (4AZR), and FLAP (2Q7M).

## Experimental procedures

### Cell culture and activation

RAW 264.7 cells (ATCC TIB71) were used for all experiments. Cells were maintained in complete media (DMEM supplemented with 5% heat-inactivated fetal bovine serum, 2 mM glutamine, 100 units/mL penicillin, and 0.1 mg/mL streptomycin). To induce PGE_2_ production, complete media was replaced with phenyl-free DMEM supplemented with 0.1% heat-inactivated fetal bovine serum and 100 ng/mL LPS (Sigma-Aldrich L5293) after washing once with PBS. Cells were incubated in LPS-supplemented media for 18 hours at 37 °C in a humidified 5% CO_2_ atmosphere. Cells were harvested by trypsin digestion for 5 min at 37 °C and centrifugation at 300 x *g*.

### PGE_2_ assay (ELISA)

Cell supernatants were removed and stored at −80 °C until use. The PGE_2_ concentration in cell supernatants was measured using a PGE_2_ Parameter Assay kit (R&D Systems KGE004B). At least 3 separate experiments were performed.

### RNA extraction and RT-PCR

Harvested cells were washed with three times using PBS. RNA was extracted from collected cells using the Qiagen RNeasy Plus Mini Kit (Qiagen) according to the manufacturer’s instructions. Reverse transcription and PCR amplification of cDNA were completed using the High Capacity cDNA Reverse Transcription Kit (Applied Biosystems) and Platinum II Hot-Start Green PCR Master Mix (Invitrogen) according to kit instructions. PCR products were visualized along with exACTGene 100 bp DNA ladder (Fisher Scientific) on a 1.5% agarose gel with 0.5 μg/mL ethidium bromide. Primer sequences used are listed in Supplemental Table 1 and PCR products were between 150 and 350 base pairs.

### Western blot

Cells were lysed in RIPA buffer with protease inhibitor cocktail (Sigma-Aldrich P8340). Protein concentration was measured using the Pierce BCA Protein Assay kit (Thermo Fisher). An equal volume of 2x Laemmli buffer (BIO-RAD) with β-mercaptoethanol was added to the cell lysate and denaturing by heating at 97 °C. Lysate (30 μg protein) per well was loaded in a 12% Mini-PROTEAN TGX gel (BIO-RAD) and then transferred to a PVDF membrane (BIO-RAD).

The membrane was incubated in blocking buffer (5% nonfat dry milk in TBST) at room temperature for 1 hour, probed overnight at 4 °C using rabbit anti-FABP5 polyclonal primary antibody (ABclonal A6373, 1:100) diluted in blocking buffer, and washed three times with TBST. The membrane was then incubated for 1 hour at room temperature in peroxidase-conjugated AffiniPure goat anti-rabbit IgG (H+L) (Jackson ImmmunoResearch AB_230739, 1:1000) diluted in blocking buffer, washed, and developed in SuperSignal West Pico PLUS Chemiluminescent Substrate (Thermo Fisher) according to the manufacturer’s instructions.

### Direct stochastic optical reconstruction microscopy (dSTORM)

Two-color dSTORM experiments were performed using a Nikon system previously described (18, 19) Approximately 300,000 cells per well were added to an 8-well chamber slide (Ibidi) and allowed to adhere for 24 hours in normal incubating conditions. The cells were then stimulated with 100 ng/mL LPS for 18 hours (as described above) or left untreated. Following 18 hr LPS activation or left untreated (control), cells were fixed and prepared for superresolution microscopy (31). Primary antibodies, anti-FABP5 (Thermo Fisher, 1:100), anti-FLAP (Novus Biologicals, 1:100), anti-COX-1 (Invitrogen, 1:100) and anti-COX-2 (Santa Cruz, 1:100) were diluted in blocking solution overnight at 4 °C. Cells were then washed twice in PBS and incubated with in-house conjugated secondary antibodies (3 μg/mL) for 60 min at RT in the dark (18, 19). For two-color dSTORM experiments, donkey anti-rabbit Alexa Fluor 647, donkey anti-goat Alexa Fluor 488 and donkey anti-mouse Atto 488 were used. The final ratio for donkey anti-rabbit:AF647 was 1:1.69 with antibody concentration 469 μg/mL. The final ratio for donkey anti-goat:AF488 was 1:2.3 with antibody concentration 97 μg/mL. The final ratio for donkey anti-mouse: ATTO488 was 1:2.5 with antibody concentration 105 μg/mL. Representative images of cells are shown and 3 separate experiments were performed.

### Clus-DoC analysis of single molecule localizations

As done previously in our laboratory (18, 19) we employed Clus-DoC (32) to quantify colocalization of individual proteins and molecules (localizations) and cluster properties in regions of interest (ROI). ROIs were drawn around the nuclear envelope and endoplasmic reticulum. Clusters were defined as having at least five localizations for FABP5, COX-1, and COX-2, and 10 for FLAP (a trimer). From 4 to 21 cells were used for analyses over 3 separate experiments.

### Time-correlated single photon counting (TCSPC) FLIM analysis

All experiments were performed at room temperature on a Nikon system equipped with Becker and Hickl DCS-120 TCSPC system (18, 19). Cells were added to an 8-well chamber slide (Millipore) and allowed to adhere for 24 hours in normal incubating conditions. The cells were then stimulated with 100 ng/mL LPS for 18 hours (as described above) or left untreated. Immediately following activation, the cells were fixed in 4% paraformaldehyde for 20 minutes at room temperature. After washing three times with PBS, the cells were permeabilized using 0.1% Triton X-100 for five minutes, washed three times with PBS, and blocked for 90 minutes in 5% donkey serum in PBS. Cells were then probed with anti-FABP5 (1:100), anti-FLAP (1:100), anti-COX-1 (1:100), or anti-COX-2 (1:100) overnight at 4 °C. The following day, cells were washed three times in PBS then incubated with Alexa Fluor 488 donkey anti-rabbit IgG and Alexa Fluor 594 donkey anti-goat IgG-conjugated secondary antibodies (1:100 in blocking buffer for 60 min in the dark. Cells were then washed twice in PBS and mounted with Vectashield containing DAPI using a 24×60-1.5 glass coverslip. Slides were sealed with nail polish and stored at 4 °C until imaged.

Protein interactions were measured by TCSPC-FLIM. The baseline lifetimes of Alexa Fluor 488 (donor fluorophore, FABP5) were calculated by single-exponential decay fitting of fluorescence emission in the absence of Alexa Fluor 594 (acceptor fluorophore, COX-1, COX-2 and FLAP). For samples stained for both donor and acceptor, lifetimes were fit to a bi-exponential decay with lifetime of one component fixed to the donor-only lifetime. The lifetime for the interacting component, **τ**_**1**_, as well as fractional contributions for the percent of interacting fluorophores, a1(%), and non-interacting component were determined. At least three separate experiments were performed, and at least 15 different pixels along the nuclear envelope were used to determine the mean value.

### Statistics

All statistics were performed using Prism Graphpad 7 (Graphpad Software, Inc.). The median and interquartile range (IQR, 25-75%) was determined. The data is expressed as: “median [25%, 75%]”. Illustrations were built in Adobe Illustrator CC (22.0.1). EIA, FLIM and cluster property results were analyzed using one-way ANOVA followed by Bonferroni multiple comparison correction or a Tukey post-hoc test, where p < 0.05 was considered significant. A non-parametric Student’s t-test with Kolmogorov-Smirnov test was performed to determine to significance where two conditions were compared (to compare medians).

## Data availability

Raw dSTORM data files are stored on a local server. dSTORM localization list text files are available on Zenodo (doi: 10.5281/zenodo.4046990). All remaining data are contained within the article.

## Funding and additional information

R.J.S. was supported by National Institutes of Health - 1R01AR065538, 1R01CA193520, R01DK062472, and S10RR027931 and the MGH Molecular Imaging Core. A.B.S. was supported by National Institutes of Health - K01DK089145 and R01DK062472. N.C.B. was supported by T32DK007540.

